# GABAA receptor mediated facilitation of proprioceptive afferent transmission to motoneurons is facilitated by TRPM8 channels on C-fibres that are gated by serotonin

**DOI:** 10.64898/2026.02.02.703408

**Authors:** Sophie Black, Yaqing Li, Krishnapriya Hari, Shihao Lin, Nicole De Silvia, Temitayo Ayantayo, Marilee J. Stephens, Keith K. Fenrich, Ana M. Lucas-Osma, Claire F. Meehan, Monica A. Gorassini, David J. Bennett

## Abstract

Sensory transmission in the monosynaptic stretch reflex (MSR) from proprioceptive Ia afferents to motoneurons is markedly inhibited by serotonin, providing a mechanism by which the brain gates spinal cord function. However, the main serotonin receptor subtype that modulates the MSR is the 5-HT1D receptor that is expressed almost exclusively on nociceptive C-fibres, suggesting an unexpected gating of proprioceptive transmission by regulation of nociceptive pathways. We demonstrate here that activation of C-fibres (electrically or pharmacologically by TRPM8 agonists) increases action potential propagation in proprioceptive Ia afferents by producing a very long-lasting tonic depolarization in Ia afferents (tonic primary afferent depolarization, tonic PAD). This tonic PAD is produced indirectly by C-fibres augmenting axoaxonic GABAergic input onto Ia afferents (bicuculline sensitive), which activates nodal GABAA receptors that aid sodium spike initiation. In contrast, activation of 5-HT1D receptors with the anti-migraine drug zolmitriptan inhibits C-fibre activity, which in turn reduces the GABAA receptor-mediated tonic PAD on Ia afferents, and reduces spike propagation to motoneurons. Overall, these findings demonstrate a mechanism by which brainstem derived serotonin not only inhibits ongoing nociceptive C-fibres activity, but also indirectly inhibits GABAergic neurons and sensory transmission in large proprioceptive sensory afferents, leading to reduced motoneuron activity. Furthermore, these findings suggest that spontaneous nociceptive C-fibres activity may provide a general increase in proprioception, opposite to the classical gating of nociceptive pathways by low threshold afferents, and impacting the clinical use of zolmitriptan.

## INTRODUCTION

The monosynaptic reflex (MSR) is produced by the activation of proprioceptive afferents that directly synapse onto motoneurons (1) and is important for the maintenance of balance and posture, forming part of a feedback loop (2–4). If the MSR becomes exaggerated, due to a lack of supraspinal inhibition following a spinal cord injury, for example, it can lead to the tremor, clonus and spasms associated with spasticity (5–7). One source of inhibitory control over the MSR is brainstem derived 5-HT (8–10). We have shown previously that the inhibitory Gi protein-coupled 5-HT_1D_ receptor potently reduces the MSR and spasticity, whereas no other 5-HT receptor types seem to directly influence sensory transmission in the MSR pathway (11). Interestingly, these 5-HT_1D_ receptors are expressed exclusively on C-fibres, and so the action of this receptor on the MSR must be indirect, perhaps inhibiting C-fibres that somehow facilitate the MSR, as previously suggested (11). Consequently, we investigated here how C-fibres and 5-HT influence sensory transmission to motoneurons in the ventral horn, with the future goal of understanding the involvement of nociceptive activity in spasticity following injury. The widespread clinical use of the 5-HT_1D_ receptor agonist zolmitriptan in treating migraines(12, 13) further motivated us in understanding how this drug influenced proprioception in the neurologically intact spinal cord.

Specialized GABAegic neurons in the spinal cord activate GABA_A_ receptors on sensory axons that lead to an axon depolarization (primary afferent depolarization; PAD) which functionally facilitates spike propagation in the axon and ultimately augments the MSR (14). This PAD arises from the unusually high intracellular chloride content of sensory neurons, compared to that of other adult neurons, and so chloride flows outward through axonal GABA_A_ receptors when they are active(14, 15). Numerous neuronal pathways activate PAD to modulate afferent transmission, including supraspinal pathways, locomotor-related circuits and a fast trisynaptic loop where afferents themselves activate excitatory neurons that innervate GABAergic neurons that recurrently activate many other afferents, usually producing a classic phasic PAD lasting about 100 ms(15–20). Contrary to long held views, PAD and associated GABAA receptors do not cause presynaptic inhibition of afferent terminal transmission, but instead GABAB receptors at the terminals cause this inhibition(14). It is now known that many of the axonal GABAA receptors that mediate PAD are located near nodes of Ranvier in myelinated proprioceptive afferents, and specifically assist in the sodium spike initiation process by depolarizing the membrane closer to spike threshold, ultimately facilitating the sensory transmission in the MSR pathway (14, 15, 21). This allows the axon to be modulated along its entire path through the dorsal columns and dorsal horn on route to the motoneurons (or brain). This nodal GABA action is particularly important at complex axon branch points in the spinal cord (and at downstream nodes), where spikes failure is more probable, especially in vivo when the spinal cord remains connected to the brain (14, 22), or when lost brainstem-derived 5-HT is supplemented by exogenous 5-HT after acute spinal transection (in vitro)(14). Whether the dependence of spike propagation on 5-HT involves 5-HT_1D_ receptors on C-fibres is unknown, a possibility we explore here.

Nociceptive C-fibres directly innervate many GABAergic neurons involved in regulation of sensory transmission, including neurons that form axoaxonic connections onto large myelinated sensory afferents, allowing them to presynaptically regulate sensory transmission (23–26). While the detailed circuits are largely unclear, such C-fibre activation produces a very long-lasting PAD in large proprioceptive afferents (tonic PAD, lasting many minutes), quite different from classic phasic PAD. This tonic PAD is partly mediated by extrasynaptic α5 GABA_A_ receptors on the proprioceptive afferents, likely activated by GABA spillover from GABAergic neurons (or astrocytes) near the terminal zones of C-fibres in the superficial dorsal horn, in a non-spiking microcircuit (8, 15, 27, 28). Thus, we explored here whether C-fibre induced tonic PAD facilities axonal conduction and sensory transmission to the motoneurons (MSR). Considering that 5-HT_1D_ receptors are exclusively on C-fibres (11) where they likely have inhibitory action (8), we also examined how these receptors inhibit the MSR. As expected, we found that activation of 5-HT_1D_ receptors with zolmitriptan inhibited C-fibres, reduced C-fibre-evoked tonic PAD and accordingly reduced sensory conduction in the MSR pathway. However, zolmitriptan did not alter the fast synaptic actions of GABA on axons in producing phasic PAD, while sharply inhibiting the MSR, suggesting that the circuits mediating tonic and phasic PAD are separate, consistent with the more general conclusion that cutaneous and proprioceptive afferents evoke PAD via separated circuits (15, 16, 20, 29).

## METHODS

Low threshold primary sensory afferents, including proprioceptive group Ia afferents were recorded to allow us to study phasic primary afferent depolarization (phasic PAD), tonic primary afferent depolarization (tonic PAD) and dorsal root reflexes (DRRs). Monosynaptic reflexes (MSRs) were evoked from dorsal root stimulation and recorded from ventral roots to allow us to see how the MSR changed in response to changes in C-fibre activity. These experiments were conducted in the whole sacrocaudal spinal cord of adult female Sprague-Dawley rats or C57BL/6 mice (2 - 5 months old) maintained in vitro or in adult awake C57BL/6 mice. All experiments were approved by the Health Sciences University of Alberta Animal Care and Use Committee.

### In vitro preparation

Under urethane anesthesia (1.8g/kg with a maximum dose of 0.45g), a laminectomy was performed and the whole spinal cord from L2 to the Ca1 region was rapidly removed and placed in oxygenated modified artificial cerebrospinal fluid (mACSF) (see (14, 15) for details). For ventral root reflex recordings, all spinal roots were removed apart from the sacral S4 and caudal Ca1 ventral roots for recording and the Ca1 or S4 dorsal roots for stimulation. For dorsal root recordings (all PAD experiments) all ventral roots were removed and S3, S4 and Ca1 dorsal roots were kept intact with usually two dorsal roots used for stimulation and four dorsal roots used for recording. For preparations involving intracellular recordings, the spinal cord was stabilised to a mesh piece of paper by gluing it ventral side down with a trace amount of cyanoacrylate. To record PAD and afferents in the deep dorsal horn, the spinal cord was glued so the left side of the cord was facing upwards to allow better access to the regions of interest. After approximately 0.5 - 1 hour in mACSF the cord was transferred to a recording chamber containing normal ACSF (nACSF) saturated with carbogen (95% O_2_ and 5% CO_2_). This chamber was maintained near 21°C with a flow rate of approximately 5ml/min.

### Intracellular recordings from afferents

To record from sensory afferents in the spinal cord, we used specialized ultra-sharp intracellular electrodes modified from those that we developed for recording motoneurons (Harvey et al. 2006; Lucas-Osma et al. 2018) to ensure that we didn’t damage the fine afferent collaterals or disturb the intracellular milieu of the afferents we were recording from. Glass capillary tubes (1.5 mm outer and 0.86 mm inner diameters with filament; 603000 A-M Systems; Sequim; USA) were pulled using a Sutter P-87 puller (Flaming Brown; Sutter Instrument, Novato, USA) to make bee stinger electrodes. These electrodes have a moderately wide final shaft (approximately 1 mm) that tapered gradually from 30 to 3 μm in length, then abruptly tapered to a final tip over the last 20 μm length. The electrodes were filled through their tips with 1 M K-acetate mixed with 1 M KCl. Our previous work found that GABAergic chloride-mediated potentials (PAD) were the same despite different concentrations of KCl, indicating that the ultra-sharp electrodes we used impeded fluid exchange between the electrode and the cell, causing no disruption to the intracellular milieu (14, 15). Following this, the electrode tips were bevelled from an initial resistance of 40 – 150 MΩ to 30 – 40 MΩ using a rotary beveller (Sutter BV-10). Electrodes were advanced into afferents using a stepper motor (666, Kopf, USA, 10 μm steps at maximal speed), typically at the boundary between the dorsal columns and the grey matter, but occasionally also deeper in the dorsal horn. All intracellular recordings were made using an Axoclamp2B amplifier (Axon Instrument and Molecular Devices, San Jose, USA) and sampled at 30 KHz (Clampex and Clampfit; Molecular Devices, San Jose, USA).

### Dorsal root stimulation, afferent identification and recording

Dorsal roots were mounted on silver-silver chloride wires above the recording chamber and covered with a 3:1 mixture of petroleum jelly and mineral oil respectively, surrounded by a high vacuum synthetic grease barrier. The dorsal roots were stimulated with a current pulse (0.1 ms) with varying intensities, expressed as a multiple of the afferent volley threshold (T) of approximately 0.003 mA. When the intracellular recording electrode was in the dorsal horn, an extracellular field corresponding to the group Ia afferent volley was observed as the first event following dorsal root stimulation. This occurred with a latency of 0.5 – 1.0 ms, depending on the root length (the roots were kept as long as possible, 10 – 20 mm), corresponding to a conduction velocity of approximately 16 – 24 m/s, as previously described for room temperature in vitro conduction(30). Our focus centred on the lowest threshold afferents, mainly proprioceptive group Ia afferents. These were identified by their direct response (spike) to dorsal root stimulation, short latency (group Ia coincident with onset of afferent volley), very low threshold (<1.5 x T) and antidromic response to ventral horn afferent terminal microstimulation (approximately 10 µA stimulation via tungsten microelectrode to activate Ia afferent terminals; tested in some afferents; discussed in more detail later in the methods). Good quality afferents used for analysis rested near – 70 mV, had a spike that peaked at about + 10 mV (overshoot) and had a brief afterhyperpolarization (10 ms). We were then able to record their electrophysiological properties, including PAD and antidromic spikes evoked by this PAD. In addition, once we penetrated and identified a Ia afferent we were able to record its membrane potential over a long time course. Once a steady baseline was achieved, we electrically stimulated C-fibres (11 stims, 1 Hz) at 50 x T and recorded the change, if any, in the Ia afferent membrane potential (see details below on selectively stimulating C-fibres). Moreover, we stimulated C-fibres chemically via the application of specific drugs to the recording chamber to activate TRP receptors found on C-fibres. Icilin is a transient receptor potential melastatin 8 (TRPM8) receptor agonist; these receptors detect menthol and cold stimuli and are activated at approximately 22°C and inactivated at approximately 26°C - 27°C (31). Zolmitriptan, as mentioned in the introduction, is a 5-HT_1D_ receptor agonist, with these receptors found exclusively on C-fibres (11, 13). We used a 3 or 10 nM dose of zolmitriptan as a low dose only activates 5-HT_1D_ receptors as opposed to higher doses which also activate 5-HT_1B_ receptors. We added icilin (10 µM) or zolmitriptan (3 or 10 nM) to the recording chamber and recorded the effects on the intracellular Ia afferent membrane potential. From our initial results we found that zolmitriptan is a useful drug to inactivate C-fibres (see results section) and therefore we were able to use this drug to determine what effect C-fibre inactivation had on several different modalities.

We also recorded from the central ends of dorsal roots cut within a few mm of their entry into the spinal cord to give the compound potential from all afferents in the root (dorsal root potential; DRP). The DRP has previously been shown to correspond to PAD, though it is attenuated compared to intracellular recordings of PAD. We found that when we wanted to record the slower components of PAD (tonic PAD) lasting many seconds or minutes, the silver-silver chloride electrodes (detailed above), sometimes had too much electrical drift at the electrode-liquid interface to be reliable. Thus we developed a miniaturized grease gap recording method to solve this problem. For this we filled one end of a 1.5 mm glass capillary tube (without filament; 628500; A-M Systems, Sequim, USA) with ACSF (or 100 µM KCl) mixed with 1.5 % agar and mounted the dorsal root onto the agar end of the tube and sealed the root in grease just above the bath (again using short roots close to the dorsal root entry zone into the spinal cord). The other end of the capillary tube was filled with 1 M NaCl and a silver-silver chloride wire was inserted into this fluid to create a highly conductive liquid-metal interface, reducing the slow drift in the electrode-liquid interface. We then recorded this signal with a DC coupled amplifier (Axoclamp2B amplifier, Axon Instruments, Molecular Devices, San Jose, USA; low pass filtered at 3 kHz) to allow us to record the DRP (tonic PAD) without extensive drift.

The rapid phasic PAD and associated evoked spikes (DRRs) were usually recorded from dorsal roots attached to the sacrocaudal spinal cord mounted on silver-silver chloride wires above the nACSF of the recording chamber and covered with a 3:1 mixture of petroleum jelly and mineral oil(30). As detailed above, it was imperative that we used very short dorsal roots cut very close to their dorsal root entry zone of the cord (within 500 µm) for this recording, so that the passive attenuation of PAD was minimized. For these experiments all ventral roots were removed and S3-Ca1 dorsal roots were kept intact with usually two dorsal roots used for stimulation and four dorsal roots used for recording purposes (standard dorsal root recording setup). This allowed us to record antidromic spikes evoked by PAD (DRRs) and phasic PAD evoked by the stimulation of one dorsal root and recorded on an adjacent dorsal root. For phasic PAD we used a custom amplifier to low pass filter the data at 3 kHz and high pass filter the data at 0.1 Hz, digitized at 30 kHz. Following dorsal root stimulation at Ia intensity (2 x T), phasic PAD arised approximately 10 ms after this stimulation with DRRs occurring on the rising phase of phasic PAD. This phasic PAD lasted approximately 100 ms. We added the drugs icilin (10 µM) or zolmitriptan (3 nM) to the spinal cord in the recording chamber to determine how C-fibre activation or inactivation affected this phasic PAD and DRRs. Using the grease gap method detailed above, we also recorded the afferent potentials (DRPs) over long periods (tonic PAD) to see how it changed with the application of icilin (10 µM), capsaicin (10 µM) and zolmitriptan (10 nM) using Axoclamp 2B amplifier (Axon Instruments, Molecular Devices, San Jose, USA; low pass filtered at 3 kHz, DC coupled without high pass filter). Capsaicin activates the transient receptor potential cation channel subfamily V member 1 (TRPV1) receptor, which detects noxious heat stimuli on C-fibres (32).

#### Ventral horn microstimulation

A method to directly assess proprioceptive afferent conduction is to selectively stimulate their terminals in the ventral horn and record the direct propagation of spikes into the dorsal roots, similar to the approaches developed by Rudomin and Wall (18, 33). We did this with a tungsten microelectrode placed in the S4 ventral horn that we stimulated minimally to reduce current spread (at ∼ 7 µA, 2 x T, 10 MΩ) and we recorded the compound action potentials that managed to propagate antidromically to the dorsal root with our standard dorsal root recording arrangement, detailed previously. From the evoked compound action potential we assessed the earliest phase to avoid later slow afferents activated by current spread. In this way, the amplitude of the compound action potential reflects the number of axons stimulated that did not fail to propagate spikes to the dorsal root. We then added zolmitriptan (3 nM) to assess its effects on the compound action potential as well as adding bicuculline (50 µM; competitive antagonist of GABA_A_ receptors) to assess the role of GABA.

#### Extracellular field recordings

Extracellular fields from the spinal cord were recorded by penetrating the spinal cord at the site of interest with sharp microelectrodes. Dorsal roots were hooked up to silver-silver chloride wires for stimulation in the main chamber to allow us to stimulate large proprioceptive sensory afferents, and we recorded the evoked extracellular field using the microelectrodes. We penetrated the cord at different locations; namely the dorsal horn and the ventral horn, and we recorded the extracellular fields evoked from the fast compound action potentials evoked in large proprioceptive sensory afferents by dorsal root stimulation (at 1 – 1.5 x T, fastest component of field). To establish the expected timing of these fields, we also directly penetrated large proprioceptive afferents in the dorsal columns to directly measure the action potential (AP; spike) evoked by dorsal root stimulation. The proprioceptive afferent field potentials had a triphasic shape: I, an initial positive phase caused by passive current driven by distal APs; II, a prominent negative phase (volley) caused by the leading edge of the AP as it reaches the recording site; III, a late positive phase (tail current) caused again by passive current but from the AP propagating past the recording electrode. It is the negative extracellular field (phase II) that allowed us to see how sensory transmission changed with C-fibre activation, as it is this component that relates to the size of the intracellular AP. As such, these extracellular field recordings can give us an indication of how well the APs are propagating down to the motoneurons in the ventral horn(14).

### Ventral root reflex recordings

The monosynaptic reflex (MSR) was recorded from ventral roots of the sacrocaudal spinal cord. Dorsal and ventral roots were mounted on silver-silver chloride wires above the nACSF of the recording chamber and covered with a 3:1 mixture of petroleum jelly and mineral oil for monopolar stimulation and recording, as we detailed previously(14, 30), though with shorter ventral roots to help record the underlying EPSP, as well as the reflexes (spikes). Ventral roots on both sides of the spinal cord were mounted for recording purposes, this included S3, S4 and Ca1 caudal ventral roots. We evoked ventral root reflexes in these sacral roots with a low threshold stimulation of the ipsilateral Ca1 or ipsilateral S4 dorsal root at T, where afferent and reflex threshold are similar (single pulses, 0.1 ms, ∼0.02 mA) using a constant current stimulator (Isoflex, Israel). The stimulation was repeated 5 times at 10 s intervals for each trail then this recording procedure was repeated every 15 minutes. The recordings were amplified (x 2000), high pass filtered at 0.1 or 100 Hz, low pass filtered at 3 kHz, and recorded with a data acquisition system sampling at 6.7 kHz (Axoscope 8, Axon Instruments). Following dorsal root stimulation an afferent volley arrived at the spinal cord approximately 0.8 – 1 ms after (due to root conduction delay), and then the MSR arose approximately 1 ms later (the synaptic delay is about 1 ms in vitro at room temperature), and therefore the MSR latency is approximately 2 ms following dorsal root stimulation. Once a steady baseline was achieved, we added icilin (10 µM) to activate C-fibres and recorded the MSR. Moreover, we stimulated C-fibres electrically by stimulating the dorsal roots at 50 x T (5 ms pulse width, detailed below) and then 10 - 20 s later we recorded the MSR evoked by a low threshold dorsal root stimulation to determine how the preceding C-fibre stimulation effected the MSR. We were confident that we were indeed recruiting C-fibres with this high threshold stimulation because the overall response increased when compared to stimulation at Ia afferent intensity (2 - 3 x T). Furthermore, we also isolated the C-fibre stimulation as detailed next.

### Selective electrical stimulation of C-fibres

We developed three methods to selectively activate C-fibres, each of which had its limitations, but we usually combined these methods to achieve just C-fibre activation. First, we maintained a very long length (∼30 mm) of a dorsal root in a bath adjacent to the main bath with the spinal cord, separated by a grease barrier and perfused with a low dose of tetrodotoxin (TTX, referred to as low TTX side bath) to the spinal cord (100 - 200 nM). This has been shown to eliminate spike generation in all sensory axons except unmyelinated C-fibres that have TTX-resistant sodium channels (28, 34, 35). We then stimulated the distal end of this dorsal root at 50 x T to activate C-fibres, to examine its effect on the MSR. In control experiments where we recorded the central end of this long dorsal root (on silver wire in grease as detailed above), we found that as the low dose TTX was applied to the side-bath all but the slow C-fibre mediated volleys were usually eliminated, confirming selective C-fibre activation, though some portion of the C-fibre volley was also usually eliminated as the dose approached 200 nM (not shown); thus a minimal dose was used that may have had small residual fast non C-fibre activation. The action of the latter on the MSR was ruled out later in experiments by a higher dose of TTX that completely blocked the root, and then stimulation had no effect on the MSR. Second, we stimulated the dorsal root in the side bath with a bipolar electrode made from gluing two fine tungsten stimulating electrodes (10 MΩ) together so that their tips were separated by only 50 µm. This arrangement readily activated the unmyelinated C-fibres, but considerably raised the threshold for activation of large myelinated afferents where the internodal spacing is near 1 mm (36), thus minimizing non C-fibre activation in the TTX side bath. The long length of the dorsal root in the side bath minimized passive current spread down the large axons into the main bath lacking TTX. Third, we developed a cathodal block circuit to stimulate C-fibres while blocking larger axons, where the applied current was raised slowly over 5 ms (with a capacitive coupling) to impose a cathodal block to nearby nodes in large afferents, and then the dorsal root was stimulated on the descending negative phase of this current, which tended to favour just slow C-fibre activation, as again confirmed in control experiments recorded from the central end of the root in grease (not shown).

In some experiments where we wanted to record from C-fibres in the dorsal root in isolation we completely blocked all spikes including C-fibre spikes with a high dose of TTX (2 µM) and eliminated fast synaptic transmission with CNQX (10 µM), a AMPA/kainite receptor antagonist; AP5 (50 µM), a NMDA receptor antagonist; strychnine (5 µM), a glycine receptor antagonist; and gabazine (10 µM), a GABA_A_ receptor antagonist. In this case the C-fibres were activated chemically as detailed above.

### Drugs and solutions

Two types of artificial modified cerebrospinal fluid (ACSF) are used during in vitro experiments. Modified artificial cerebrospinal fluid (mACSF) was used in the dissection dish and the normal artificial cerebrospinal fluid (nACSF) was used in the recording chamber. The mACSF was composed of (in mM) 118 NaCl, 24 NaHCO_3_, 1.5 CaCl_2_, 3 KCl, 5 MgCl_2_, 1.4 NaH_2_PO_4_, 1.3 MgSO_4_, 25 D-glucose, and 1 kynurenic acid. The nACSF was composed of (in mM) 122 NaCl, 24 NaHCO_3_, 2.5 CaCl_2_, 3 KCl, 1 MgCl_2_, and 12 D-glucose. Both types of ACSF were saturated with carbogen (95% O_2_ and 5% CO_2_) and maintained at a pH of 7.4. The drugs added to the nACSF were capsaicin, icilin, gabazine, CNQX, AP5, bicuculline (all from Tocris, Minneapolis, USA) and TTX (TRC, Toronto, Canada). All drugs were dissolved as a 10 – 50 mM stock in distilled water before final dilution in nACSF and applied to the spinal cord in vitro. The drugs zolmitriptan and strychnine (both from Tocris, Minneapolis, USA) were first dissolved in a minimal amount of DMSO (final concentration in nACSF of 0.02 %; by itself DMSO has no effect on in vitro DRP, DRRs or MSRs in vehicle controls) in order for the drug to fully dissolve, which was then added to the nACSF and added to the spinal cord in the recording chamber.

### Genetically labelling GAD2 neurons

We have bred a strain of mice with GAD2 neurons, a specific subpopulation of GABAergic neurons, labelled with a fluorescent reporter; enhanced green fluorescent protein (EGFP), to visualize GAD2 neurons. To do this, GAD-cre-ER mice are crossed with reporter mice with floxed stop codons on sequences for EGFP (flx-EGFP mice). The mice are given tamoxifen to induce cre (2 x 4 mg intraperitoneal) at 4 weeks old(14).

### Horizontal ladder walking

We examined mouse hindlimb placement accuracy on a horizontal ladder with 2 mm brass rungs with 2 cm spacing between rungs. Mice were naïve to the ladder walking (untrained). The ladder was elevated ∼20 cm above the ground with one end resting on their home cage so that they felt relaxed and motived walking to their cage. They were filmed at 240 frames/s (with an iPhone15 camera, Apple, USA), which allowed quantification of rapid slips. Hind foot placing errors were counted, and quantified as a proportion of the number of attempts (usually 20 - 30 attempts, as mice traversed about that many rungs). An error was defined as an outright miss of the rung with the foot falling below the ladder, or a poor placement (on tip of toes) that led to a slip off the rung. This procedure was carried out in mice injected with zolmitriptan (i.p. 20 mg/kg in a vehicle of 3 ml saline with 2% DMSO) or a vehicle control, with mice restrained in a small tube immediately after the injection so that they did not practice walking prior to testing, and then tested on the ladder at 12 min after the injection, to allow the drug to reach peak effect when tested.

### Immunolabelling

Transgenic mice, outlined above, were euthanized with Euthanyl (BimedaMTC; 700 mg/kg) and perfused intracardially with 100 ml of saline containing sodium nitrite (1 g/l; Fisher) and heparin (300 IU/l, from 1,000 U/ml stock; Leo Pharma) for 3 - 4 minutes, followed by 400 ml of 4% paraformaldehyde (PFA; in phosphate buffer at room temperature), over 15 minutes. Spinal cords were postfixed in PFA overnight at 4°C and then immersed in 30 % sucrose in phosphate buffer for 72 hours. The spinal cord tissue was coated with O.C.T cryoprotectant (Sakura Finetek) before being frozen in 2-methylbutane (Fisher Scientific) and then cut on a cryostat NX70 (Fisher Scientific) in transverse 25 µm sections. The tissue was stored at -20°C until histological processing. We mounted the spinal cord sections on slides and rinsed with Tris-buffered saline (TBS, 50 mM) containing 0.3% Triton X-100 (TBS-TX, 10 minute rinses used for all TBS-TX rinses). Next, nonspecific binding was blocked with a 1 hour incubation in 10 % normal donkey serum (NDS; ab7475, Abcam) in PBS-TX. The primary antibodies: sheep anti-calcitonin gene-related peptide (CGRP; 1:5000; ab22560, Abcam), mouse anti-Bassoon (1:400; SAP7F407, ENZO) and rabbit anti GFP (1:500; A11122, Life Technology), were diluted in 2 % NDS in PBS-TX and applied to slides overnight at room temperature. The following day, the tissue was rinsed with PBS-TX (3 × 10 mins) and incubated with fluorescent secondary antibodies: donkey anti-sheep Alexa Fluor 647 (1:500; ab150179, Abcam), donkey anti-mouse Alexa Fluor 555 (1:500; ab150106, Abcam) and donkey anti-rabbit Alexa Fluor 488 (1:500; A-21206, Thermofisher) diluted in 2% NDS in PBS-TX for 2 hours at room temperature. After rinsing with PBS-TX (2 × 10 mins) and PBS (2 × 10 mins), the slides were covered with Fluoromount-G (00-4958-02, ThermoFisher Scientific, Waltham, USA) and coverslipped in Permount (Sakura Finetek USA, Torrance, CA, USA).

Standard negative controls in which the primary antibody with either 1) omitted or 2) blocked with its antigen (quenching) were used to confirm the selectively of the antibody staining, and no specific staining was observed in these controls. For antibody quenching, the peptides used to generate the antibodies (AAP34984, Aviva Systems Biology, San Diego, USA) were mixed with the antibodies at a 10:1 ratio and incubated for 20 hours at 4°C. This mixture was then used instead of the antibody in the above staining procedure.

### Image acquisition

Image acquisition was performed with a Leica TCS SP8 confocal Laser Scanning Microscope (Leica Microsystems Inc.). Leica Application Suite X (Leica Microsystems CMS GmbH) was used for visualization and analysis of all images taken.

### Data analysis

To effectively quantify PAD we measured the compound effects of PAD by measuring the dorsal root potential (DRP) from the cut end of dorsal roots to allow us to record steady DC potentials without extensive drift (Figs 1B-C; 4B- F; 5A). When the DR was stimulated it evoked a phasic PAD and associated DRP, the latter an order of magnitude smaller in absolute potential than the average phasic PAD recorded intracellularly (14). We determined the effects of drugs on the afferent membrane potential (tonic PAD) by normalizing the changes in DRP with the drugs by the ratio of the phasic DRP for each root to the average size of intracellular phasic PAD, the latter which is approximately 3.7 mV(15). Data was analyzed using Clampfit 8.0 (Axon Instruments, USA) and Sigmaplot (Jandel Scientific, USA) and is shown as a mean ± standard deviation (SD, the latter used to quantify variability). A paired *t*-test was used to test for statistical differences, with a significance level of *P* < 0.05.

**Figure 1.**
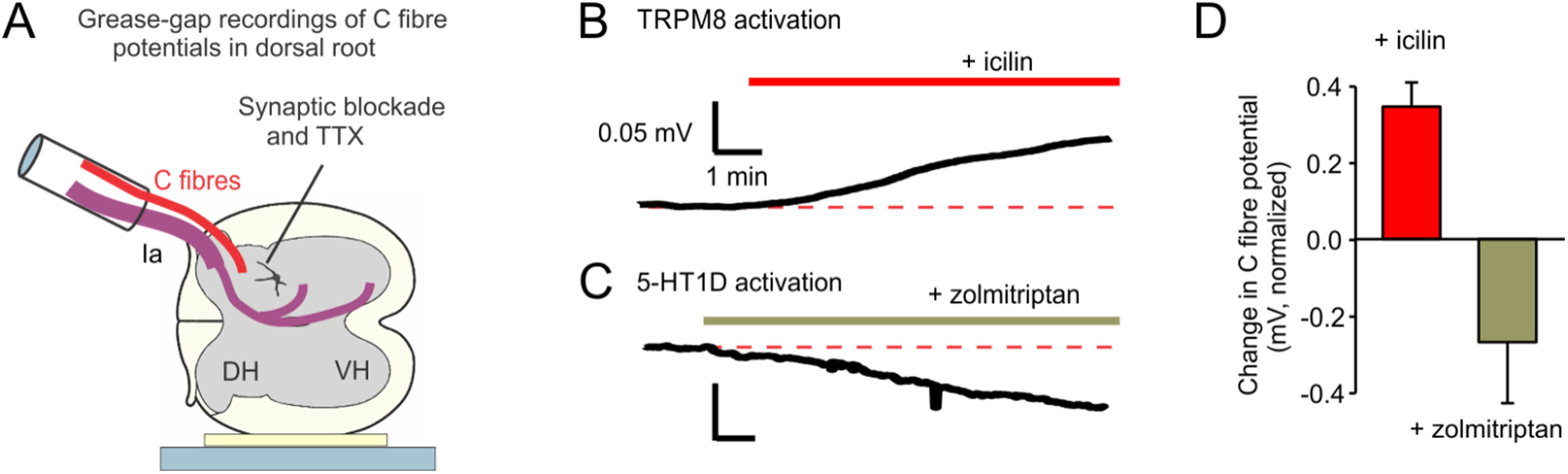
Icilin depolarizes C-fibres whereas zolmitriptan hyperpolarizes C-fibres, demonstrating that these drugs are useful tools to either activate or inactivate C-fibres. **A:** Recording of C-fibre potential changes in dorsal roots in the presence of TTX (2 µM) and full synaptic block (10 µM CNQX, 50 µM APV, 5 µM strychnine and 10 µM gabazine) to block all spike transmission in the whole adult rat spinal cord, maintained in vitro. **B:** Application of the TRPM8 receptor agonist icilin (10 µM) depolarized C-fibres in the presence of full synaptic block. **C:** Activation of 5-HT_1D_ receptors via zolmitriptan (3 nM) hyperpolarized C-fibres in the presence of full synaptic block. **D:** Following the application of icilin to the spinal cord the C-fibres depolarized (n = 5), whereas following the application of zolmitriptan (n = 5) the C-fibres hyperpolarized in the presence of TTX and full synaptic block, shown as the change in compound membrane potential relative to control, normalized by ratio of DR root evoked phasic DRP to expected phasic PAD (3.7 mV; see Methods). Error bars SD. * significantly different, *P* < 0.05.

## RESULTS

### C-fibres are activated by icilin and inhibited by zolmitriptan

To initially study C-fibres in isolation we bathed the whole spinal cord in TTX (2 µM) and synaptic transmission blockers (10 µM CNQX, 50 µM AP5, 5 µM strychnine, 10 µM gabazine), while recording the central cut ends of dorsal roots in the whole isolated in vitro adult rat spinal cord. When we then activated C-fibres using the TRPM8 agonist icilin (10 µM; Andersson et al. 2004; Mergler et al. 2007), the dorsal roots depolarized (dorsal root potential, DRP; Fig 1B), indicating an overall depolarization of C-fibres since these TRP receptors are exclusively found on C-fibres (Richter et al. 2019; De- Schepper et al. 2008; Lucas). In contrast, the selective 5-HT_1D_ receptor agonist zolmitriptan (3 nM; Tepper et al. 2002; Thomsen et al. 1996) hyperpolarized the sensory axons (DRP; Fig 1C). Because these 5-HT receptors are also found exclusively on C-fibres (11, 13), this indicates that the C-fibres were hyperpolarized by zolmitriptan (consistent with the inhibitory actions of 5-HT_1D_ receptors in other neurons, including activation of GIRK potassium channels). Overall, these results demonstrate that zolmitriptan and TRP agonists like icilin are effective tools for inactivating or activating C-fibre actions, respectively, which we used extensively throughout our experiments.

#### C-fibre activity increases the MSR to motoneurons

We next examined whether C-fibres and 5-HT modulated sensory transmission to motoneurons by recording the monosynaptic reflexes (MSRs) from the ventral roots in response to dorsal root stimulation at proprioceptive sensory afferent intensity (2 x T). For this, we selectively stimulated C-fibres by taking advantage of the relative resistance to TTX of the sodium channels in C-fibres, compared to sodium channels of other afferents. Specifically, we stimulated the C-fibres in the distal portion of a dorsal root isolated by a grease barrier from the rest of the cord and bathed in TTX (abbreviated TTX side-bath; see Methods). When C-fibres were selectively stimulated in this way, the MSR recorded 20 s later was strongly facilitated (Fig 2B), without altering motoneuron activity at these long delays (i.e. no postsynaptic facilitation lasting 10-20 s). This MSR facilitation built up with repeated C-fibre stimulation (at 20 s or 3 s intervals) and lasted several minutes after stimulation ceased (Fig 2E). Likewise, selective activation of C-fibres with icilin (10 µM) facilitated the MSR (Fig 2C), again without altering motor output prior to MSR testing.

**Figure 2.**
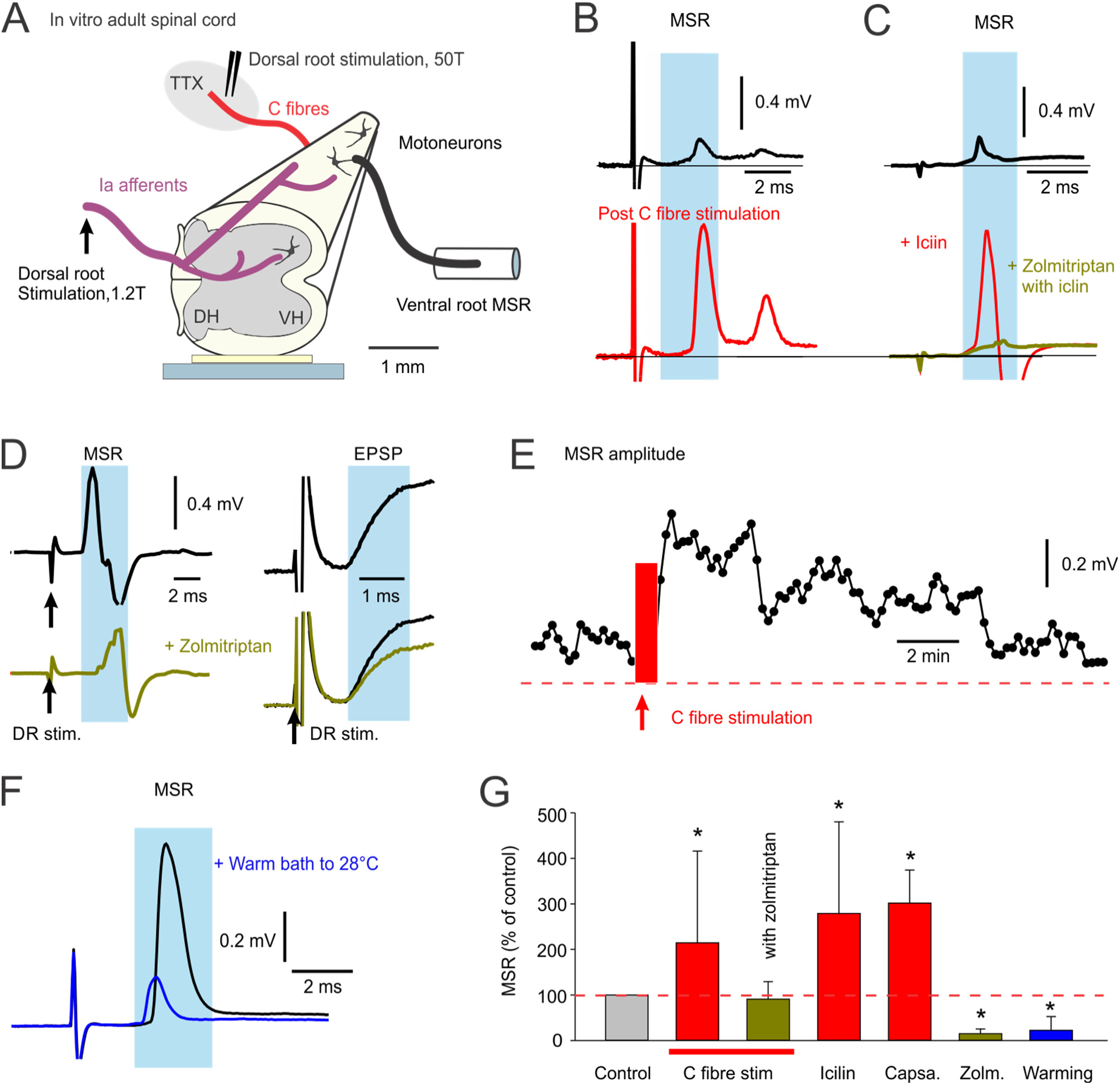
Monosynaptic reflexes (MSRs) are facilitated by C-fibre activation and inhibited by C-fibre inactivation. **A**: Schematic of whole adult in vitro rat spinal cord. The MSR was recorded from an S4 ventral root evoked from a dorsal root stimulation (Ca1 DR, 1.2 x T, 0.1 ms pulse, T sensory threshold), which was sometimes preceded by an isolated stimulation of C-fibres in the S4 DR (with DR isolated by a grease barrier in a side bath containing low dose TTX; 100 – 200 nM, 0.5 - 1 ms pulse at 50xT, bipolar stimulation). **B**: The MSR was measured repeatedly every 20 s prior to any C-fibre stimulation, and then again 20 s after a C-fibre stimulation (50 x T, 1 ms pulse, at 20 s intervals), repeated 3 times at 20 s intervals (averages of 3 trials shown). Ventral root activity was not different with and without C-fibre stimulation at the time of the MSR test (n = 19/19). **C**: Activation of C-fibres with the TRPM8 agonist icilin (10 µM) also facilitated the MSR, and again did not change motoneuron activity (VR activity, n = 11/11), whereas the application of zolmitriptan (3 nM) after icilin reduced the MSR (n = 5/5 similar). **D**: Zolmitriptan (3 nM) reduced the MSR and associated monosynaptic EPSP (the latter subthreshold to spiking). **E**: Brief repeated C-fibre stimulation (11 stims, 0.3 Hz, 50 x T) facilitated the MSR for several minutes (n = 19/19 similar). **F**: MSR decreased when the bath temperature was raised to 28°C to inactivate TRPM8 receptors. **G**: On average, C-fibre stimulation (n = 19), icilin (n = 11) or capsaicin (10 uM, n = 3) increased the MSR, zolmitriptan blocked this C-fibre action (n = 6), and either zolmitriptan alone (n = 6) or warming the bath from 22°C to 28°C (n = 5), reduced the MSR. MSR shown as a percentage of control MSR. Mean ± SD. * significantly different, *P* < 0.05

In contrast, the application of zolmitriptan markedly reduced the MSR and associated EPSP (Fig 2D), and blocked the facilitation of the MSR by icilin or C-fibre stimulation (Fig 2C and G). As we have previously reported, zolmitriptan has no direct action on motoneurons (37), suggesting again that it acts on the sensory axons and not motoneurons. Overall, this suggests that C-fibres somehow affect proprioceptive afferent transmission to motoneurons, and even do so spontaneously, prior to inhibiting C-fibres with zolmitriptan.

Icilin activates TRPM8 receptors which respond to cool temperatures, activating near room temperature (around 22°C) where we normally maintain our in vitro bath, and off at higher temperatures.

Interestingly, when we warmed the spinal cord to 28°C to turn off the inhibitory action of TRPM8 receptors on C-fibrers, we found that the MSR was strongly inhibited (Fig 2F), again consistent with there being an ongoing tonic C-fibre activity that facilitates the MSR, as shown by an increase in the MSR with icilin, though other temperature effects (38) may confound temperature change experiment.

### C-fibre activity facilitates sensory transmission to the ventral horn

To confirm that C-fibre activation increases proprioceptive afferent transmission to the ventral horn, we directly measured from these afferents while manipulating C-fibre activity (Fig 3A). Using sharp microelectrodes we penetrated the cord in the dorsal horn and the ventral horn, and we recorded extracellular fields from action potentials evoked in large Ia proprioceptive sensory afferents by low threshold dorsal root stimulation (at 1 – 1.2 x T, fastest component of field corresponding to afferent volley; Fig 3B). To determine how these fields relate to the afferent action potential, we also made intracellular recordings from the large branches of these proprioceptive afferents in the dorsal horn (Fig 3B), as previously detailed (14). The earliest part of the extracellular field represents the afferent volley from these large afferents, and had a distinctive shape with at least two phases, followed by more complex synaptic fields: I, an initial positive phase caused by passive current driven by distal action potentials approaching the electrode; and II, a prominent negative phase (volley) caused by inward sodium current when the action potential reached the recording site (Fig 3B-C; as in (14, 39). The relative size of the extracellular field (phase II) allowed us to estimate how sensory transmission changed with C-fibre activation, as it reflects the compound effect of the action potentials reaching the electrode from multiple nearby afferents.

**Figure 3.**
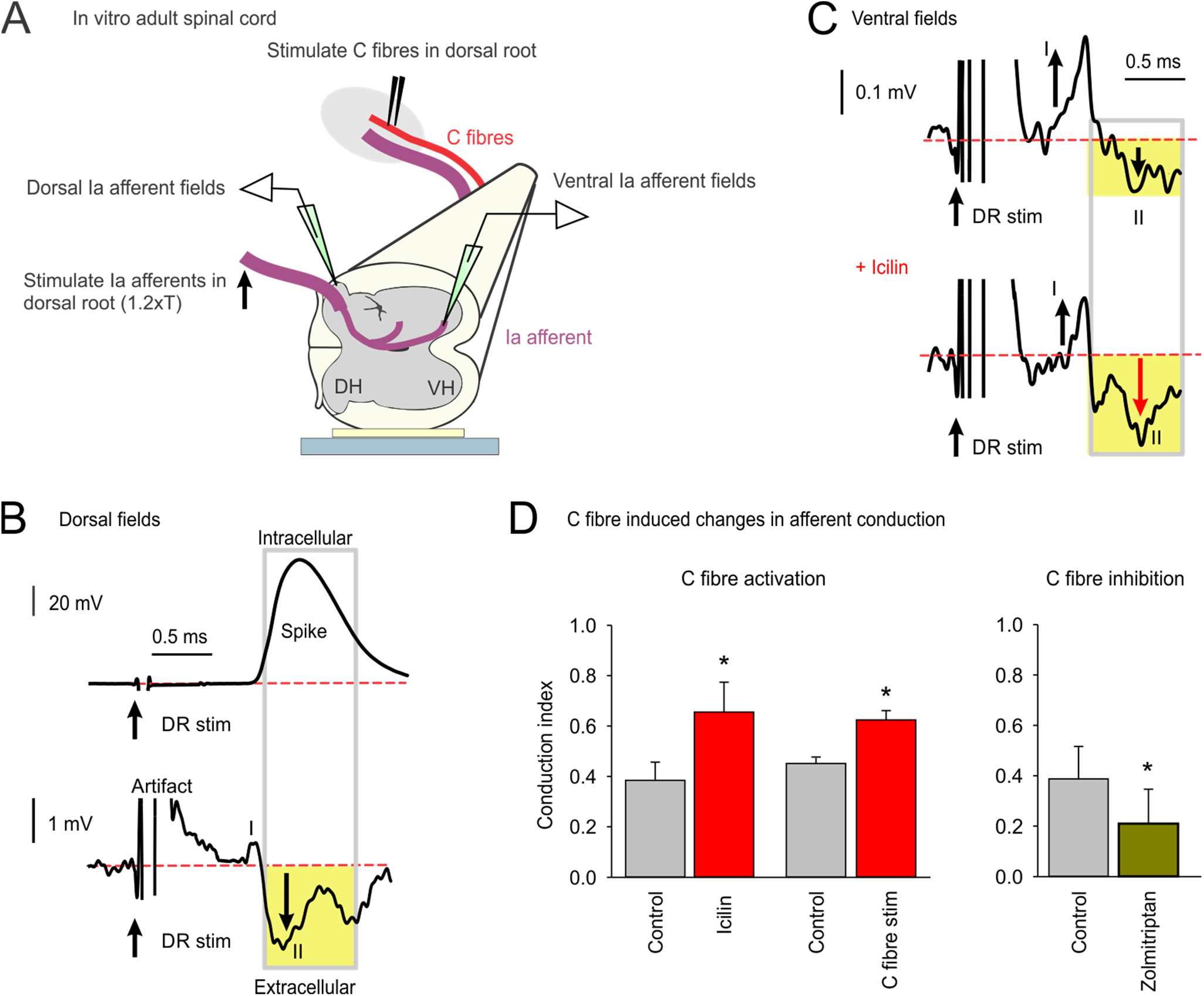
Activation of C-fibres increases sensory transmission to the ventral horn. **A**: Schematic of recording afferent fields in adult in vitro spinal cord. **B**: Extracellularly recorded field potential (bottom) from compound action potentials evoked in large proprioceptive afferents by dorsal root stimulation (at 1.2 x T, 0.1 ms pulse; S4 or Ca1 root, T sensory threshold), compared to action potential (spike) recorded intracellularly by penetrating a nearby afferent (bottom; n = 20/20 similar). Averages of stimulation at 2 Hz. Recordings from dorsal horn near dorsal columns. **C**: Extracellular fields evoked by same dorsal root stimulation (1.2 x T), but recorded in the ventral horn at the Ia afferent terminals (terminal potential fields). The fields had at least a biphasic shape: I, an initial positive phase caused by passive current driven by distal action potentials (spikes); and II, a prominent negative phase (volley, yellow) caused by the leading edge of the spikes as they reach the recording site; after this the field was complex and involved synaptic events and so ignored. Selective C-fibre stimulation with icilin (10 µM) increased the proprioceptive afferent negative volley (phase II). Averages of > 60 recordings over several minutes. **D**: To quantify these actions of C-fibres on the terminal field in the ventral horn, we used the positive field (*pf, phase I*) and negative field (*nf, phase II*) fields of the triphasic shape of the afferent terminal extracellular field to compute a *conduction index = nf / (nf+pf)*. Following the activation of C-fibres by or icilin (n = 5) or selective C-fibre stimulation (field measured 1 – 20 s after adjacent DR stimulation 50 x T, 11 x 5 ms pulses at 1 Hz, with root bathed in TTX or CNQX and APV, as in Methods, n = 4) the conduction index increased, whereas inhibition of C-fibres with zolmitriptan (0.1 µM, n = 6) decreased the conduction index. Mean ± SD. * significantly different, *P* < 0.05.

When we measured this extracellular at the afferent terminals in the ventral horn of the spinal cord (Fig 3 C) the negative phase of the fields was relatively smaller (II), and instead the initial positive phase (I) was more prominent, consistent with many dorsally generated axon spikes passively depolarizing the ventral axon terminal, but failing to propagate spikes to the terminal, as previously detailed(14). We quantified afferent conduction to the ventral horn by normalizing the amplitude of the negative phase of the field (phase II, denoted nf) by the total peak-to-peak size of the field from the positive peak (in phase I, denoted pf) to the negative peak (nf + pf) to compute a Conduction Index (CI) = nf / (nf + pf). Prior studies have shown that failing spikes lose the negative phase II volley, but produce a larger positive phase I, from the passive effect of spikes at upstream to the failure point (14, 39), giving some indication of the total spikes prior to the failure point. This CI provided us with an index approximating the proportion of non-failing spikes: where a CI of 1.0 indicates no failure (pf = 0) and 0.0 indicates complete failure (pf >> nf). Being a ratio, this index also compensates for the large variability in the absolute afferent volley size between animals and recording location.

Activating C-fibres via the application of icilin (10 µM; Fig 3C) increased the ventral Ia afferent volley (increased phase II, and decreased phase I) and overall increased the conductance index, consistent with increased conduction. Similarly, brief selective C-fibre stimulation (for 11 s, as detailed above) increased the Ia afferent volley (phase II) for several minutes (Fig 3D). In contrast, silencing C-fibres with zolmitriptan decreased this volley and the conduction index (phase II, Fig 3D). Together these results suggest that C-fibre activity increases spike propagation to proprioceptive afferent terminals near motoneurons (likely by preventing conduction failure, see Introduction).

#### C-fibre activation tonically depolarizes large proprioceptive afferents by activating GABAergic neurons that activate α5 GABAA receptors on these afferents

We next examined how C-fibres increase afferent transmission, by recording intracellularly from large myelinated proprioceptive afferents. Selective C-fibre activation (50 x T, with root bathed in TTX, as detailed in Methods) elicited a long-lasting depolarization of large proprioceptive afferents (tonic PAD, Fig 4A-B), which was partly blocked by the extrasynaptic α5 GABA_A_ receptor antagonist L655708 (Fig 4B,D), indicating that C-fibres act indirectly by activating GABAergic innervation of afferents to produce tonic PAD. Tonic PAD has previously been shown to increase afferent conduction (14, 15), and so these results are consistent with our findings that C-fibres increase afferent transmission to motoneurons (Figs 2 – 3). In contrast, when C-fibres were inhibited by zolmitriptan the proprioceptive afferents were hyperpolarized (Fig 4E), demonstrating that the C-fibres acted upon by zolmitriptan must have been spontaneously active, steadily facilitating tonic PAD in proprioceptive afferents, likely via activation of GABAergic neurons that activate the GABA receptors on these afferents.

**Figure 4.**
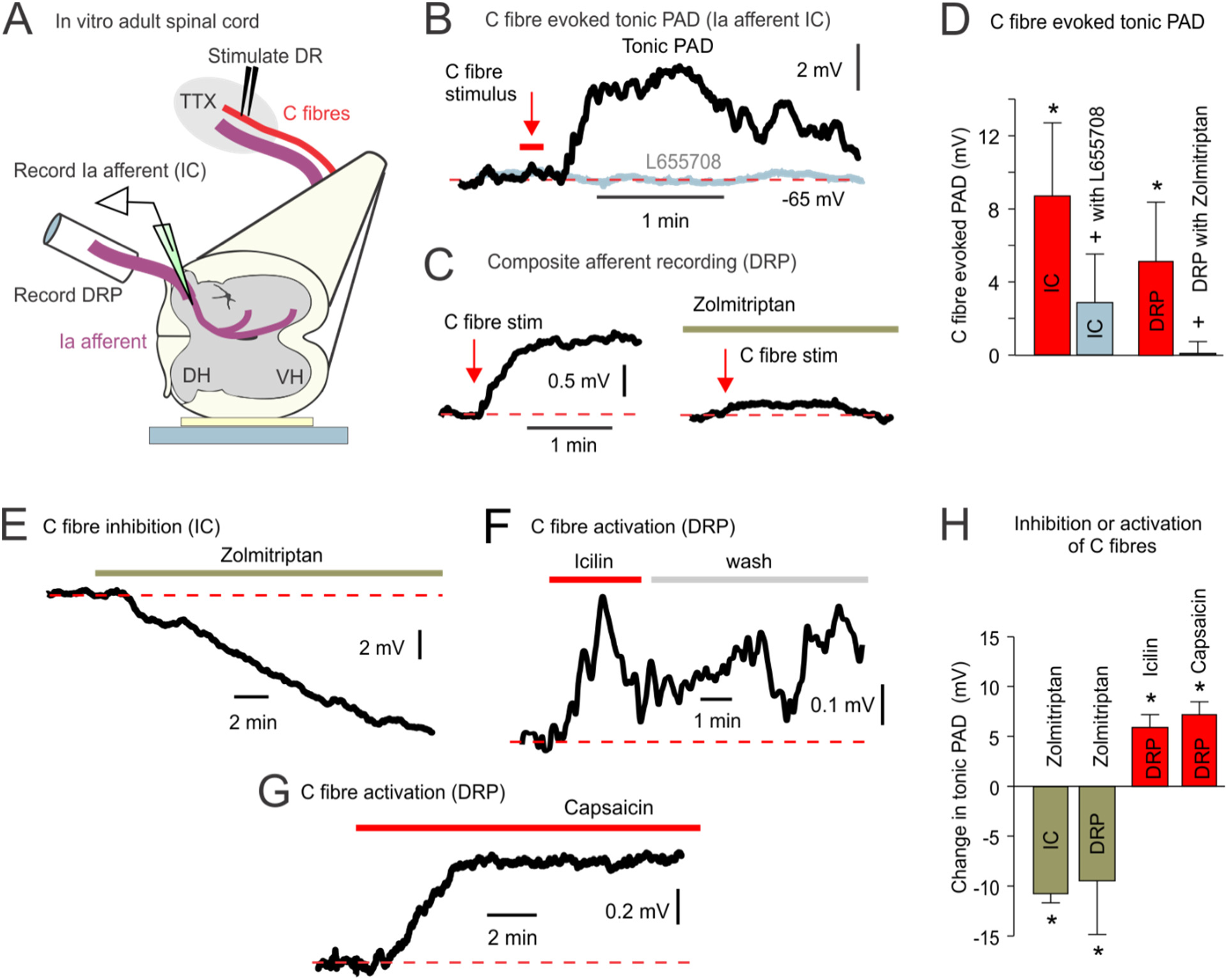
Activation or inhibition of C-fibres increase or decreases tonic PAD in proprioceptive afferents. **A**: Intracellular recording from proprioceptive group Ia afferent (S4 afferent) in the dorsal horn, and composite recording from sensory axons in the dorsal root (DRP), both with isolated C-fibre stimulation (applied to dorsal root, in 200 nM TTX, 2 or 11 pulses of 0.5 ms at 0.2 - 1 Hz, 50 x T). **B**: C-fibre stimulation (11 pulses at 1 Hz) depolarized the Ia afferent for several minutes (tonic PAD), but L655708 (0.1 uM) blocked this. Data filtered at 1 Hz. **C**: C-fibre stimulation (2 pulses at 0.2 Hz) also depolarized dorsal root for minutes (compound tonic PAD) and zolmitriptan (3 uM) blocked this. **D**: Group averages of C-fibre induced depolarization of Ia afferent (IC, n = 6) and dorsal root (DRP; n = 5), and reduction of this depolarization with L655708 (IC, n = 5) or zolmitriptan (DRP, 3 nM, n = 5). * significant increase, + significant reduction in C-fibre action, p < 0.05. Mean ± SD. **F:** Intracellular recording from a proprioceptive Ia afferent (S4 afferent) in the dorsal horn, with zolmitriptan (3 nM) causing a hyperpolarization (n = 4/4 similar). **F-G**: Icilin or capsaicin (10 µM) depolarized dorsal roots (DRP) for many minutes (tonic PAD) (n = 5 and 3 respectively). **G**: Group averages of the hyperpolarization of Ia afferents (IC, n = 4) or dorsal roots (DRP, n = 5) by zolmitriptan, and depolarization of dorsal roots by icilin (n = 6) or capsaicin (n = 3). * significantly change, *P* < 0.05.

To further quantify the modulation of PAD by C-fibres, we also recorded the composite PAD from many afferents on the central cut end of dorsal roots (DRP), which mainly represents the PAD from large myelinated afferents (including proprioceptive Ia afferents(14)). Again, selective C-fibre activation tonically depolarized the afferents (Fig 4C, F-H), regardless of whether we selectively activated C-fibres electrically (in TTX side bath) or chemically with the TRP channel agonists icilin or capsaicin, consistent with depolarization of large proprioceptive afferents (tonic PAD). This C-fibre induced tonic PAD was eliminated by prior application of zolmitriptan to silence C-fibres (Fig 4D), consistent with the 5-HT_1D_ receptor inhibitory action on C-fibres. Also, C-fibre inactivation with zolmitriptan again hyperpolarized the sensory axons (Fig 4H).

This dorsal root recording approach to study overall afferent depolarization (DRP) is possibly contaminated by direct contributions from C-fibres in the recording. However, the bulk of the DRP evoked by C-fibre stimulation is likely due to depolarization of large diameter afferents (like proprioceptors) for several reasons. First, this DRP far outlasted the C-fibre stimulation (Fig 4B, C, F) and expected transient depolarization of the C-fibres themselves. Second, the much smaller size of C-fibres likely made any direct depolarization of C-fibres an order of magnitude smaller than the depolarization observed in large proprioceptive afferents (compare Figs 1 and 4), due to much greater electrotonic attenuation of centrally generated voltages in small vs large axons (Hari et al 2022), which we recorded a few mm away on the dorsal root. Third, 5-HT_1D_ receptors are only in the portion of C-fibres in the dorsal horn (Lucas 2019) and thus only hyperpolarize C-fibres electronically far from where we record on the root, again giving much smaller effects compared to the indirect hyperpolarization of Ia afferents by zolmitriptan (compare Fig 1C to Fig 4E). Finally, a blockade of GABA receptors with L655708 prevented most of the C-fibre mediated depolarization of large proprioceptive afferents (Fig 4 B,D), indicating that the indirect action of GABA on these large afferents causes the main depolarization in the dorsal root.

#### C-fibre activation increases antidromic spike propagation on proprioceptive afferents

The mechanism by which this C-fibre-induced tonic PAD facilitates afferent transmission (MSR) likely involves facilitation of sodium spike conduction in the branch points of afferents, far from afferent terminals(14, 15), as we have seen with indirect extracellular recording of terminal field potentials near motoneurons (Fig 2). One way to further test the idea that this PAD aids spike propagation is to examine antidromically propagating spikes that are readily observed when a phasic PAD evoked by stimulation of Ia afferents (in an adjacent dorsal roots) is large enough to directly evoke a spike. Thus, using both direct intracellular recordings from Ia proprioceptive afferents and dorsal root recordings, we measured such PAD-evoked antidromic spikes. As discussed previously(15), while these antidromic spikes are often generated by a fast rising phasic PAD (about 80% of the time), they only rarely fully propagate to the dorsal root (9% of the time produce a spike, DRR), likely due to the difficulty of spikes propagating backwards from small axons branches into larger parent branches(40). Thus, during intracellular recordings from Ia afferents we commonly observed small partial spikes evoked by phasic PAD (failure potentials, FPs; Hari), which represent failed spikes and their the passively signal from the last active node prior to spike failure, as shown in Fig 5A-B. However, when we selectively stimulated C-fibres 20 s prior to the Ia stimulation, this improved the antidromic spike evoked by phasic PAD induced from the Ia stimulation, with a full action potential propagating to the electrode (Fig 5B-C). Likewise, when we again stimulated Ia afferents in the dorsal roots to evoke a phasic PAD, we observed antidromically propagated spikes on adjacent roots (dorsal root reflexes, DRRs; Fig 6A, D) that were facilitated by the application of icilin (Fig 6D). Overall, this is consistent with the notion that the C-fibres induced depolarization of Ia proprioceptive afferents helps with the initiation of spikes by phasic PAD.

**Figure 5.**
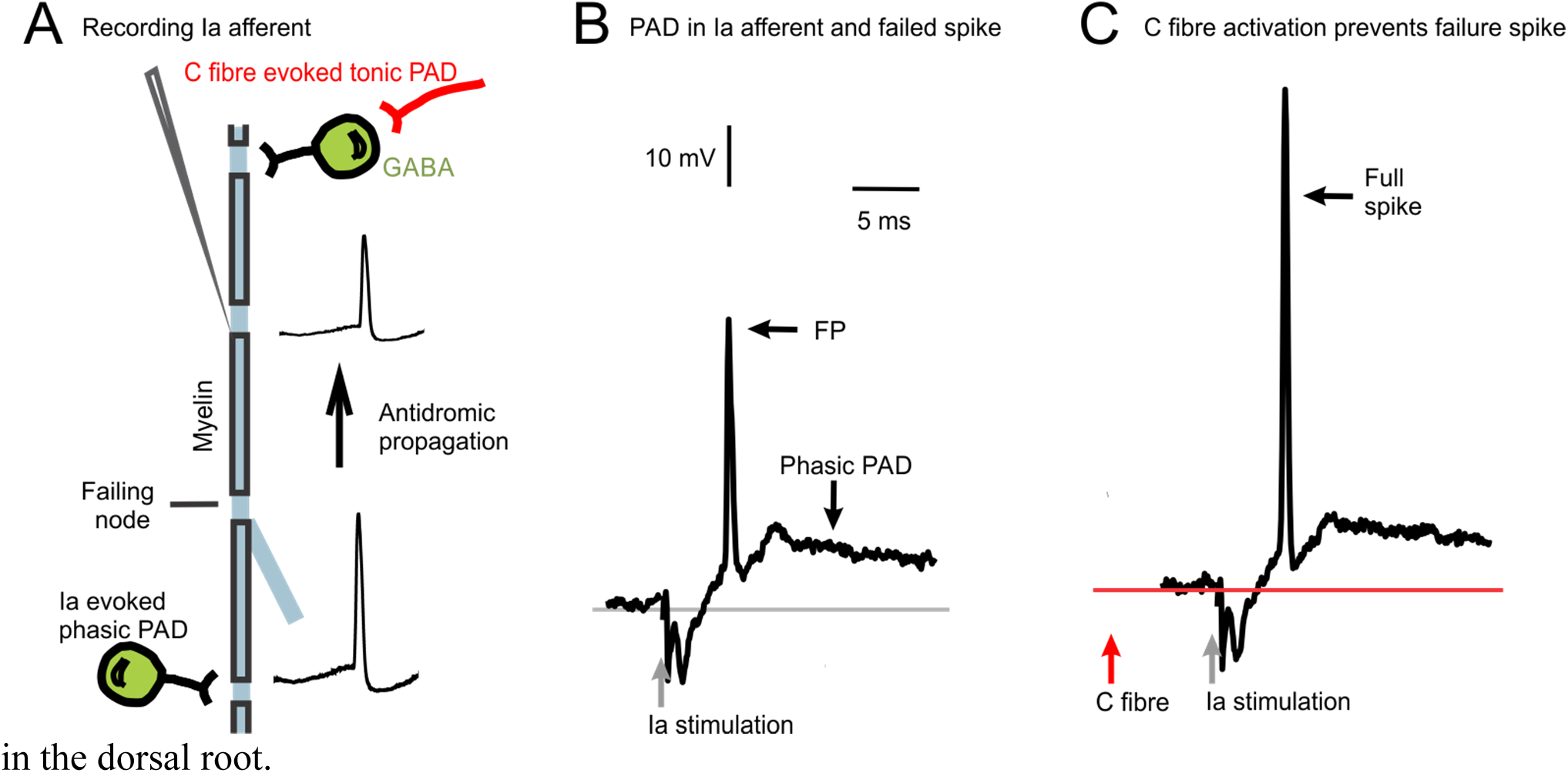
Activation of C-fibres prevents failure of spikes in primary afferents. **A**: Intracellular recording from group Ia afferent in dorsal horn, and suggested locations of GABAergic input (green cells) that produce PAD from Ia afferent and C-fibre stimulation. **B:** Low threshold dorsal root stimulation to activate Ia afferents (1.1 x T, 0.1 ms) evoked a large phasic PAD (Ia evoked) that produced a partial spike that failed to conduct antidromically to the electrode (FP). **C**: Selective C-fibre stimulation (50 x T, 5 ms pulse, in TTX side bath) tonically depolarized the afferent (tonic C-fibre evoked PAD) and 20 s later improved the phasic PAD-induced antidromic spike in the same afferent (n = 5/5 afferents similar).

**Figure 6.**
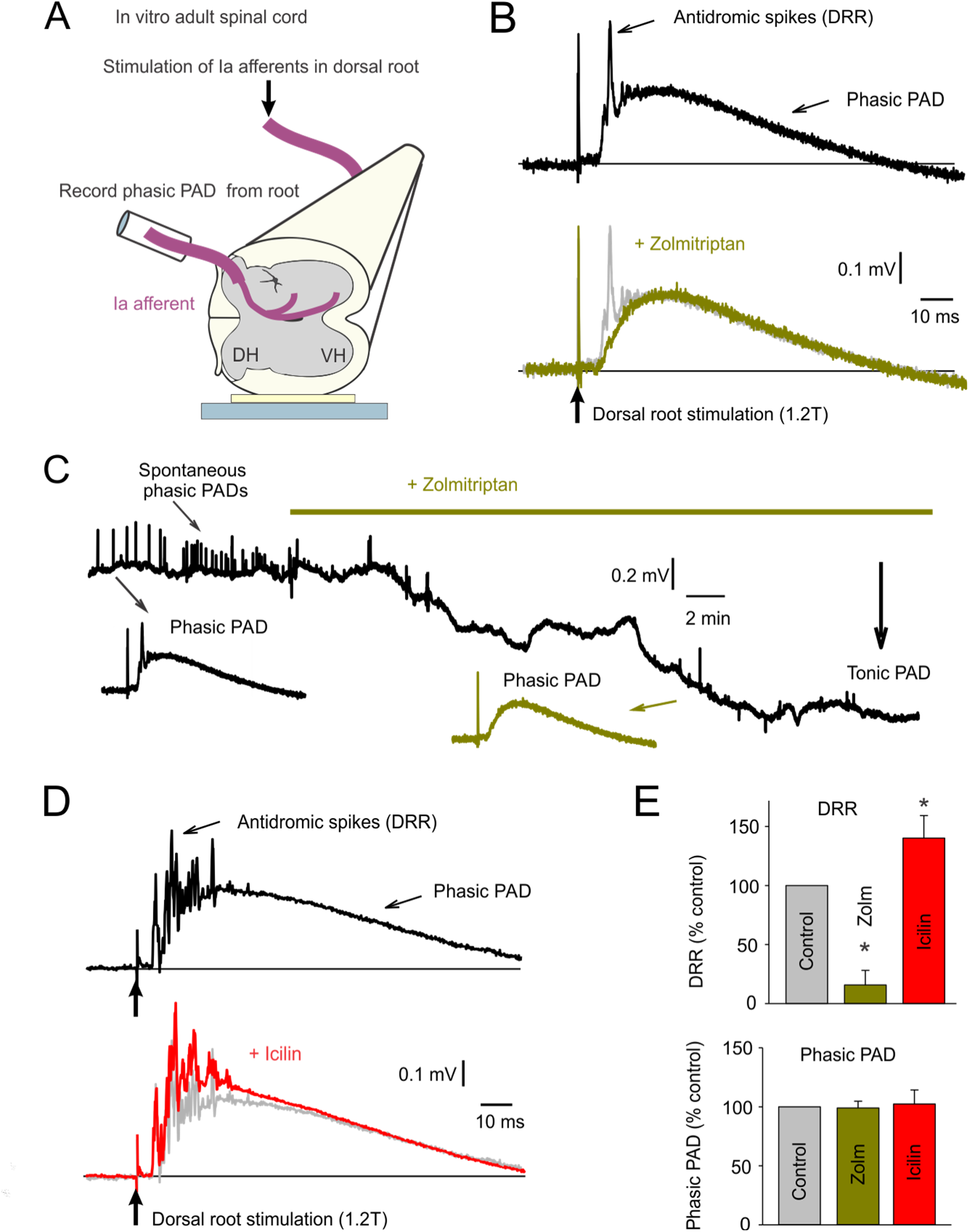
Inhibiting C-fibres with zolmitriptan hyperpolarizes proprioceptive afferents and decreases antidromic spike transmission, whereas activating C-fibres has the opposite effect. **A**: Schematic of setup to record PAD and DRR from the cut end of a dorsal root in response to stimulation of an adjacent dorsal root in whole adult rat spinal cord maintained in vitro. **B**: Following low intensity dorsal root stimulation (1.2 x T, 0.1 ms pulse, group I stimulation) a phasic PAD occurred. This phasic PAD evoked antidromic afferent spikes (DRRs) on its rising phase (black trace). Zolmitriptan (10 nM) did not change phasic PAD, but reduced the antidromic spikes (DRRs, green; pre-zolmitriptan trace overlayed in grey). **C**: Change in PAD with time after application of zolmitriptan (10 nM), with a tonic hyperpolarization and loss of spontaneously evoked PADs (small irregular depolarizations), but not loss of dorsal root evoked phasic PAD (insets from same root as B). **D**: Similar phasic PAD to B, but icilin (10 µM) increased the antidromic spikes (DRRs). **E**: Group averages of action of zolmitriptan (n = 8) and icilin (n = 5) on the DRR and phasic PAD. Mean ± SD. * significantly change from control, *P* < 0.05.

In contrast, when we inactivated C-fibres with zolmitriptan, and accordingly hyperpolarized the Ia afferents, the antidromic spikes evoked by phasic PAD were reduced (Fig 6B-E, DRR decreased), even though dorsal root evoked phasic PAD itself did not change (Fig 6E). As mentioned in the Introduction, phasic PAD is the classic PAD which has been associated with presynaptic inhibition, though it is not causative, with GABAB instead causing presynaptic inhibition. Therefore, these results suggest again that decreased C-fibre activity (via zolmitriptan) reduces Ia afferent conduction, seen as reduced DRRs, whereas phasic PAD and presynaptic inhibition play no role. The reduced DRRs likely occurred as a result of zolmitriptan hyperpolarizing the afferents (reducing tonic PAD) and brought some afferents below spike threshold to propagate backwards along the axon. Chronologically, these observations of zolmitriptan decreasing afferent conduction were the initial impetus for us thinking that PAD assisted sodium channels and led to our recent thorough analysis of this idea(14), and we only now understand and report these zolmitriptan results with hindsight. Interestingly, there were spontaneously occurring phasic PADs (left of Fig 6C) that were eliminated by zolmitriptan, indicating that spontaneous synaptic inputs to the afferents were somehow also reduced by silencing C-fibres, even though Ia evoked phasic PAD was not, suggesting that these two sources of PAD are independent.

A method to more selectively assess proprioceptive afferent conduction is to stimulate their terminals in the ventral horn next to motoneurons and record the direct propagation of their spikes into the dorsal roots(18, 33) (Fig 7A). Because only large proprioceptive afferents innervate the motor nucleus (especially Ia afferents), this method provides a way to selectively target these sensory axons. Using a microelectrode placed in the S4 ventral horn we stimulated minimally to reduce current spread (at ∼ 7 µA) and recorded the compound action potentials that managed to propagate antidromically to the dorsal root. We also assessed the earliest phase of the compound action potential to avoid later slow afferents activated by current spread. In this way the amplitude of the compound action potential reflects the number of Ia (or possibly group Ib) axons stimulated that propagate spikes to the dorsal root. When we applied zolmitriptan to block C-fibres, this compound action potential recorded on the dorsal root was markedly reduced (Fig 7B-C), and importantly this action of zolmitriptan was blocked by inhibiting GABA_A_ receptors with a pre-treatment of bicuculline (Fig 7C), suggesting that antidromic conduction depended on C-fibre activity with an obligatory involvement of GABA neurons. Blocking GABA_A_ receptors alone mimicked the action of zolmitriptan (Fig 7D), again reducing the compound action potentials, consistent with a large part of the action of GABA on afferents depends on C-fibre activity.

**Figure 7.**
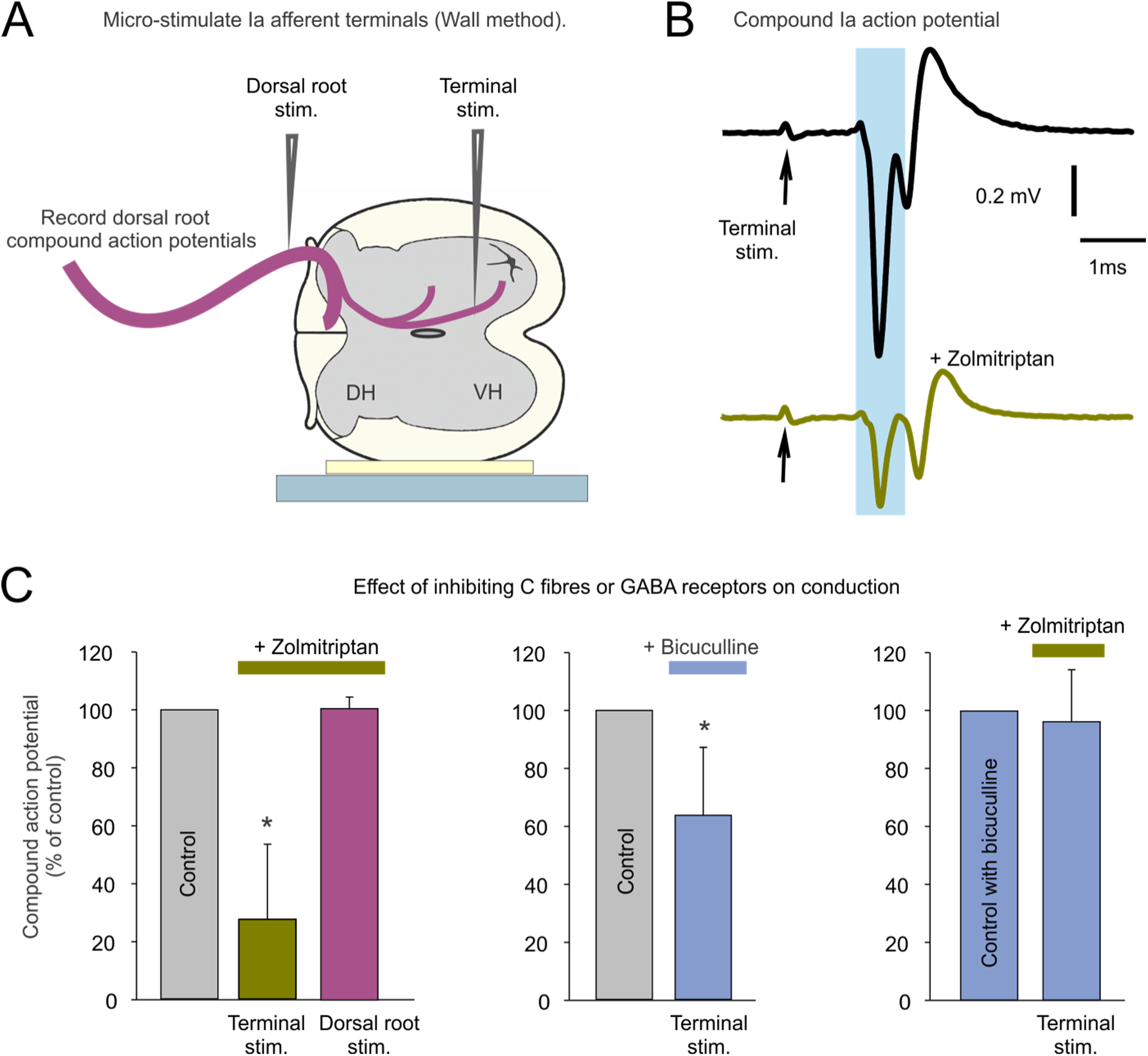
Inhibiting C-fibres with zolmitriptan reduces sensory transmission to and from the ventral horn. **A**: Micro-stimulation of Ia afferent terminals in the ventral horn in the isolated spinal cord (1.5 x T, 0.5 ms pulse), and recording of antidromically propagated compound action potentials recorded from dorsal roots. **B**: Zolmitriptan (10 nM) reduced the Ia compound action potential (n = 7). **C**: Zolmitriptan (3 nM) decreased the compound action potential (n = 7), but following inhibition of GABA_A_ receptors with bicuculline or gabazine (50 µM or 5 µM, combined with 50 uM APV, 50 uM CNQX and 5 uM strychnine) the compound action potential was no longer decreased by zolmitriptan (n = 8). Bicuculline (50 µM) alone decreased the compound action potential, mimicking the action of zolmitriptan. Mean ± SD. * significantly change from control, *P* < 0.05.

Stimulating the dorsal root, rather than the terminals, produced compound action potentials that were unaffected by zolmitriptan (Fig 7C), consistent with the lack of 5-HT1D receptors on the roots and a central action of this receptor on C-fibres. Taken together these results are again consistent with our conclusion that inhibition of C-fibres reduces an endogenous tonic GABA tone (GABA-mediated tonic PAD) that normally facilitates afferent spike initiation and conduction.

#### C-fibres make direct contacts with GAD2 neurons

To determine how C-fibres activate the GABAergic neurons underlying PAD, we next examined the relation of dorsally located C-fibres to GABAergic neurons. We immunolabelled C-fibres with CGRP (Fig 8) and their presynaptic terminals with Bassoon (Fig 8). GAD2^+^ neurons, a subpopulation of GABAergic neurons known to form axoaxonic connections and produce PAD in large myelinated axons, were genetically labelled in GAD2//EYFP^+^ mice (Fig 8). We found that C-fibres directly synapse onto GAD2 neurons (Fig 8; arrows, with Bassoon on C-fibres presynaptic to the GABAergic neuron). This establishing an anatomical basis for C-fibre mediated PAD in proprioceptive afferents, though other GABAergic neurons or astrocytes cannot be ruled out, especially considering how long-lasting C-fibre actions are.

**Figure 8.**
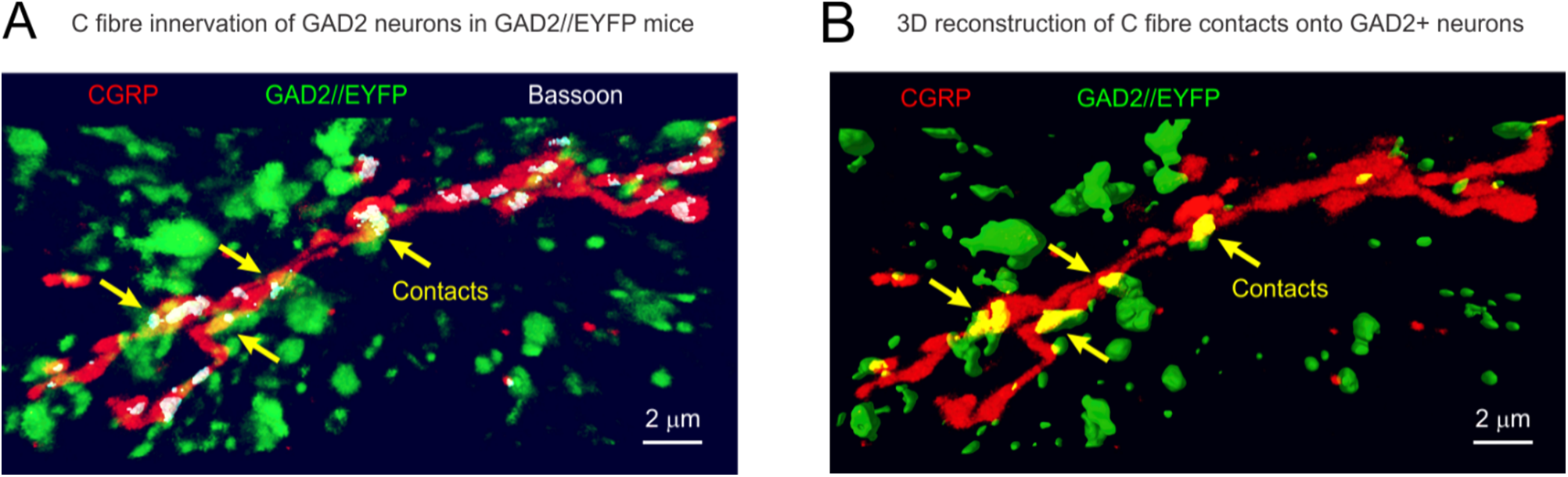
C-fibres innervate GABAergic neurons. **A**: C-fibres immunolabelled with CGRP make direct contacts onto genetically labelled GABAergic GAD2^+^ neurons (in GAD2//EYFP mice) in the dorsal horn (n = 5 similar). Presynaptic contacts of C-fibres are immunolabelled with axonal bassoon, with only the bassoon in the 3D reconstructed volume of the C-fibre shown, for clarity. **B**: 3D reconstruction of GAD2^+^ neurons and their innervation by C-fibres, with contact zones labelled in yellow, and indicated with arrows.

#### Zolmitriptan increases proprioceptive errors on horizontal ladder walking

Considering that proprioceptive sensory transmission to motoneurons is impaired, we examined whether zolmitriptan caused a more general loss of proprioceptive transmission that affects limb movements. For this we quantified foot placement errors on a horizontal ladder where proprioception is needed to place the hind feet that are not visible, quantified by counting the proportion of times a foot missed or slipped off a ladder rung. Naïve mice not training on the ladder made errors in 18.7 ± 11.4% of foot placement attempts, whereas these errors increased to 43.0 ± 9.7% when zolmitriptan was administered (i.p; P < 0.05, mean ± SD, t-test, n = 18 and 4, respectively), consistent with a similar loss of proprioception that we see in the reflex pathways.

## DISCUSSION

Our results demonstrate that spike propagation in group Ia proprioceptive sensory axons and associated MSR transmission are facilitated by activation of nociceptive C-fibres, including icilin- and temperature-sensitive C-fibres (TRPM8), capsaicin-sensitive C-fibres (TRPV) and TTX-resistant C-fibres (Na1.8 expressing and stimulated 50xT, including most C-fibres). This facilitation of spike propagation by C-fibres is indirect, involving activation GABAA receptors on the myelinated axon branches in the spinal cord that cause a tonic depolarization of these axons (C-fibre PAD), because it is sensitive of GABA receptor blockers. As recently reported by Hari et al. (14), phasic PAD evoked by direct or indirect activation of GABAergic neurons that innervate nodes of sensory axons aids spike propagation by producing a subthreshold depolarization that assists sodium spike initiation at the many complex branch points that are otherwise prone to conduction failure, but generally does not cause a large enough depolarization to induce suprathreshold sodium channel inactivation (conduction block). The C-fibre evoked tonic PAD that we report here likely acts similarly to facilitate spike propagation in myelinated sensory axons, because C-fibre PAD directly aids nodal spike initiation by bringing axons closer to threshold, mimics the facilitation of spikes by optogenetic activation GABAergic neurons that innervate sensory axon nodes(14), and is reduced by GABAA receptor antagonists or 5-HT receptor agonists that inhibit sensory axon conduction. Unexpectedly, we also find that Ia afferent conduction under resting conditions depends on spontaneous activity of C-fibres and GABA receptors, since conduction is sharpy reduced by either inhibiting C-fibre activity with zolmitriptan or blocking GABAA receptors, without any prior C-fibre stimulation, which has important functional implications that we discuss further below.

### Neuronal circuits that mediated nociceptive facilitation of PAD and sensory transmission

The neuronal circuits that mediate C-fibre facilitation of proprioceptive afferent sensory transmission are currently only poorly defined, though we know some critical elements. C-fibres directly innervate the many GABAergic neurons of the dorsal horn (41), including GAD2+ neurons (Fig 9), and in general GABAergic neurons provide the dorsal horn with a dense plexis of GABAergic terminals, with many more GABAergic terminals than in the ventral horn(41–43). We have also recently shown that large proprioceptive afferents have extrasynaptic and synaptic GABA receptors along their whole length as they traverse through the dorsal horn on the way to the ventral horn, and it is these dorsally located GABA receptors that produce both tonic and phasic PAD (not terminal GABA_A_ receptors; (14, 15)).

Thus, it is reasonable to suppose that these dorsally located extrasynaptic GABA receptors on afferents are strongly activated by spillover of GABA from the dense GABAergic neuron innervation of the spinal cord. A caveat to this argument is that repeated optogenetic activation of GAD2^+^ neurons does not produce a long-lasting facilitation of proprioceptive afferent transmission comparable to the prominent after-effects of repeated sensory axon stimulation(14). This suggests that another source of GABA also contributes to axon conduction, possibly including astrocytes that release GABA(44), or GAD2^−^GABAergic neurons. Both Hounsgaard’s group(28) and our group(15) have shown that spillover of GABA from C-fibre stimulation does depolarize large proprioceptive afferents (producing tonic PAD), and this is mediated via extrasynaptic GABA receptors (L655708 sensitive; (15)). This PAD can persist even when spikes are blocked in all but C-fibres (which have TTX-resistant sodium channels), and thus must at least in part be mediated by a local non-spiking circuit in the superficial dorsal laminae where the C-fibres terminate (15, 28), again consistent with astrocyte action. Thus, the substrate by which C-fibres facilitate large proprioceptive afferent transmission at a minimum involves a simple dorsally located circuit where C-fibres activate GABAergic astrocytes or neurons which, in turn, activate extrasynaptic GABA receptors on these large afferents. We show that C-fibres make direct contacts with GAD2+ neurons, a unique genetically defined GABAergic neuron population that make contacts with afferent nodes and terminals (14, 43), though again GAD2 neurons alone are unlikely to provide the bulk of GABA that modulates tonic PAD (14, 15). Likely other more complex pathways are also involved (14), including C-fibres activating other neurons or astrocytes, which could release other transmitters like glutamate, in addition to GABA, onto afferents (20, 28, 29, 45). Evidently the pathways that underly phasic and tonic PAD differ, because C-fibres can increase tonic PAD without changing phasic PAD, both phasic and tonic PAD can independently change afferent transmission.

### Inhibition of the MSR pathway by GABAB and not GABAA receptors

While various subpopulations of GABAergic neurons release GABA onto nodes (14) and terminals of proprioceptive afferents (14, 15, 42, 46), the afferent terminals innervating motoneurons appear to mostly express GABA_B_ receptors, rather than GABA_A_ receptors, and these GABA_B_ receptors are primarily responsible for presynaptic inhibition of sensory transmission to motoneurons (sensitive to GABA_B_ blockers), contrary to the long-standing belief that GABA_A_ and PAD are involved (14, 47). In contrast, the axonal GABA_A_ receptors that tend to mostly be near nodes are too far from ventral terminals to be involved in presynaptic inhibition. However, these GABA_A_ receptors can have a paradoxical action, that has historically been confused with presynaptic inhibition, as follows. When axonal GABA_A_ receptors are synaptically activated by GABAeric neurons they produce a fast phasic PAD that can be large enough to directly evoke spikes in the afferents. This phasic PAD is particularly easy to evoke experimentally, even in humans, since a large synchronous volley evoked in sensory axons themselves activates first order glutamatergic neurons that broadly activates the GABAergic neurons that produce phasic PAD in widely distributed sensory axons, in a well known trisynaptic loop (16, 29, 48). The spikes evoked in proprioceptive afferents by such phasic PAD travel orthodromically to the motoneurons and ultimately activate the MSR pathway, including producing monosynaptic EPSPs in antagonists muscles that normally only receive reciprocal inhibition from the stimulated sensory nerve (14, 15, 48). Paradoxically, these indirect monosynaptic EPSPs evoked by phasic PAD depresses the excitability of the MSR pathway to subsequent activation, due to post-activation depression of the synaptic machinery, explaining why PAD has previously been mistaken for presynaptic inhibition (48). Ironically, a clear example of such indirect monosynaptic EPSPs is seen in Figure 6 of the original 1959 publication of Frank that led to the study of presynaptic inhibition, though it was not noticed at the time (49).

PAD-evoked spikes sometimes also travel antidromically out of the dorsal root, making them readily detectable as a dorsal root reflexes (DRRs), which serves as proxy for the likely presence of post-activation depression (15, 18, 47, 48). We demonstrate here that decreasing C-fibre activity (with zolmitriptan) markedly decreases these phasic PAD related spikes (DRR), by simply hyperpolarizing the axons, making spike initiation less likely. This finding further supports the role of C-fibres in promoting spike propagation. It also suggests that C-fibre activity assists in post-activation depression of the MSR associated with PAD-evoked spikes, and this would normally be inhibited by brainstem-derived 5-HT that inhibits C-fibre activity, explaining why phasic PAD most readily facilitates, rather than inhibits, the MSR in the intact spinal cord (in vivo vs in vitro)(14). In contrast, decreasing C-fibre activity (with zolmitrptan) had little effect on phasic PAD itself, further emphasizing the conclusion that the circuits mediating tonic and phasic PAD are separate (15, 16, 20, 29).

### Functional consequences of nociceptive PAD

The increased proprioceptive afferent conduction produced by C-fibres has numerous important functional consequences related to both modulation of complex spinal motor circuits and sensory transmission to the brain. We have detailed here the simplest of these functions, the MSR connecting afferents to motoneurons, which underlies the stretch reflex. Our results demonstrate that the increased sensory conduction produced by C-fibres and GABAergic activity leads to increased EPSPs and MSRs on motoneurons. Perhaps the most surprising finding is that even spontaneous C-fibre activity facilitates sensory transmission to motoneurons (increased MSR), since the MSR is reduced by reducing C-fibre activity (with the 5-HT_1D_ agonist zolmitriptan) in the absence of exogenous activation of C-fibres.

Indirect in vivo measurements suggest similar conclusions, because 5-HT_1D_ agonists (zolmitriptan or sumatriptan) likewise inhibit the MSR in vivo (9, 50). This suggests that C-fibres are spontaneously active enough to contribute to a basal facilitation of sensory conduction. C-fibres are peculiar because they express their sensory receptors (TRP receptors) all along their length in the spinal cord, as well as in the peripheral tissue they innervate. Thus, local events in the spinal cord (temperature, pH, etc.) contribute to overall C-fibre activity, and combined with peripheral inputs, C-fibres are known to exhibit a degree of spontaneous activity (51–54). The TRPM8 receptor (which detects cold and menthol stimuli; activated by icilin) is particularly interesting in this regard because it is activated at room temperature(31), and thus in the periphery (in vivo) is spontaneously active. Thus, something as simple as skin temperature appears to increase sensory transmission and reflexes in the spinal cord.

Since nociceptive C-fibre activity increases afferent transmission throughout the spinal cord, it stands to reason that C-fibre activity should increase sensory transmission up the dorsal columns, raising the intriguing possibility of increased acuity of proprioceptive sensation with nociceptive activity. If this is the case, it might be a useful adaptation, providing high sensory acuity when responses to pain are required. Whether noxious input actually does increase proprioceptive perception has proven difficult to study, because guarding effects induced by pain change motor responses and distraction induced by pain confound such studies. Direct recordings from the ascending dorsal column pathway are needed to settle this issue. Our find that zolmitriptan inhibits C-fibre activity and decreases proprioceptive reflexes, raises the intriguing possibility that zolmitriptan use in treating migraines might impair proprioception, perhaps explaining the tripping and clumsy movements anecdotally reported with such triptan use (Claire Meehan, unpublished results), as we see with mice on a horizontal ladder.

### Serotonergic inhibition of sensory transmission

The present study was initially motivated by our finding that 5-HT_1D_ receptors are located exclusively on C-fibres, and not alpha motoneurons or large afferents, even though they strongly inhibit the MSR(11). Thus, the question arose as to how these 5-HT receptors can modulate the MSR without being located on any element of the MSR pathway. We initially explored the actions of C-fibres on PAD, naively thinking C-fibres should increase PAD and thereby decrease the MSR, since PAD has been classically associated with presynaptic inhibition of the MSR (18, 33, 47). However, we found the opposite. C-fibre activation produces a tonic PAD (15). We now know that this tonic PAD facilities afferent conduction, rather the causing presynaptic inhibition(14). This solves the problem of how serotonin receptors inhibit the MSR: brainstem derived 5-HT activates 5-HT_1D_ receptors that inhibit C-fibre activity which, in turn, reduces GABAergic neuron (or glial) activity and extrasynaptic GABA receptor activity on proprioceptor afferent nodes, which in turn reduces spike conduction on the afferent (likely via an increase in branch point failure) and this ultimately reduces the MSR. This rather convoluted mechanism of the action of 5-HT_1D_ receptors likely explains some anomalies in the action of the 5-HT_1D_ agonist zolmitriptan, which have previously been reported. For example, after the application of zolmitriptan to the spinal cord, the resulting reduction in the MSR is difficult to reverse by washing zolmitriptan (11), which likely involves the 5-HT_1_ receptor somehow irreversibly reducing the spontaneous TRP receptor activity on the C-fibres. This is plausible considering the intimate coupling of 5-HT_1_ receptors and TRPM8 (55). Also, the finding that knockout of 5-HT_1D_ receptors leads to an increased MSR can now be viewed as mediated by possible increased C-fibre activity (56).

### Functional implications and relation to pain gating

In summary, we show the existence of a long-lasting C-fibre-induced tonic PAD which increases sensory transmission to motoneurons. This occurs via a dorsally located microcircuit activating GABAergic neurons or astrocytes which, in turn, activate extrasynaptic GABA_A_ receptors located at nodes, likely reducing branch point failure. When C-fibres are not active, this tonic PAD and associated facilitation of spike transmission ceases. Functionally, this C-fibre and associated GABAergic activity leads to increased sensory transmission to motoneurons, and may well more generally leads to increased sensory transmission to the brain, as well as to complex spinal motor circuits. This C-fibre mediated facilitation of non-noxious sensory transmission in large proprioceptive afferents stands in contrast to classic concepts of gating of sensory transmission (57–59), and thus warrants further study, especially as it relates to pain.

## ACKNOWLEDGMENTS

We thank Leo Sanelli for technical assistance. Research was supported by Canadian Institute of Health Research (CIHR, MOP14697 and PJT169029) and the National Institutes of Health (NIH, R01NS47567).

## Notes

### Competing Interest Statement

The authors have declared no competing interest.

### Summary of Updates

We have added an author we omitted. We have also added a paragraph to the end of the Results section. No conceptual changes were made to the paper.

